# Position-dependent changes in phycobilisome abundance in multicellular cyanobacterial filaments revealed by Raman spectral analysis

**DOI:** 10.1101/2023.02.14.528583

**Authors:** Jun-ichi Ishihara, Yuto Imai

## Abstract

The one-dimensional filamentous cyanobacterium, *Anabaena* sp. PCC 7120, shows a simple pattern consisting of two types of cells under the nitrogen-deprived conditions. We found that the microbial pigment composition in differentiated (heterocyst) and undifferentiated cells (vegetative cells) were distinguished by using a Raman microscope. The Raman bands of phycocyanin and allophycocyanin were higher in the vegetative cells than those in the heterocysts. However, these bands were statistically lower in a part of vegetative cells, which were located far from a nearby heterocyst. That is, the pigment composition in the individual cells was affected by a locational information in a filament.

## Description

The filamentous and multicellular cyanobacterium, *Anabaena* sp. PCC 7120 (hereafter named *Anabaena*), is composed of a lot of vegetative cells connected in a one-dimensional manner (Figure A). The vegetative cells are specialized for photosynthesis and contain both PSI (Photosystem I) and PSII (Photosystem II). Under nitrogen-deprived conditions, *Anabaena* differentiates into cells named heterocyst, which are specialized for nitrogen fixation, at an interval of approximately ten cells along the filament [1–7]. Once a heterocyst is fully differentiated, it never reverts to a vegetative cell. As the number of vegetative cells increases by the cell division, a new heterocyst is differentiated approximately midway between two existing heterocysts. A heterocyst is easily distinguished by using a standard microscope as it is larger in size and rounder in shape. In addition, the light-harvesting phycobilisome complex is chemically decomposed or inactivated in heterocysts, leading to suppress the oxygen-evolving PSII [6–9]. The heterocysts and vegetative cells exchange metabolites produced by the nitrogen fixation or photosynthesis with the neighboring cells because the photosynthesis and nitrogen fixation are incompatible in the same cell [4–7] (Figure A).

The differences of the photosynthetic systems between vegetative cells and heterocysts have been frequently studied by using a fluorescence microscope [8–12]. Recently, we analyzed microbial pigment compositions in the vegetative cells and heterocysts in a non-invasive and non-labeling manner by a Raman spectral measurement [13, 14]. Fifteen vegetative cells (or heterocysts) were selected randomly from several filaments, and the Raman spectra were measured for every single cell. The Raman bands in the spectra were assigned to vibrational modes of resonance Raman bands of light-harvesting pigments including chlorophyll *a*, phycocyanin, and allophycocyanin. Especially, in the heterocysts, the intensities of Raman bands of phycocyanin and allophycocyanin were remarkably decreased when compared to those of chlorophyll *a*. Given that phycocyanin and allophycocyanin are components of the light-harvesting phycobilisome complex in PSII, our previous result shows good correspondence with the previous studies that phycobilisome is decomposed during the differentiation (a chemical decomposition reduces the amount of the target molecule and the corresponding Raman intensity) [8, 9].

The aim of this study is to investigate how the pigment compositions in the individual cells were affected from a nearby heterocyst. To address, we measured Raman spectra of individual cells along a filament of *Anabaena* by selecting the central points of the cells in a sequential manner. The average Raman spectra of vegetative cells and heterocysts, which consist of a whole filament (Filament 1), were shown in Figures C and D. The procedure to obtain the Raman spectrum is explained in Materials and Methods section. The intensity values of a Raman spectrum in the region of 990 to 1770 cm^-1^ were normalized to unity, and the normalized spectra of all vegetative cells (or heterocysts) in the filament were averaged (Figures C and D). The band positions in the Raman spectra of the vegetative cells were nearly the same as those of the heterocysts. Hereafter, the normalized band intensities at 1328, 1629, and 1639 cm^-1^ were selectively used as the signal of chlorophyll *a*, phycocyanin, and allophycocyanin, respectively [14]. To show the difference of spectral features more clearly, the Raman spectrum in heterocysts was subtracted from that in vegetative cells (Figure E). Consequently, the intensities of Raman bands at 1629 and 1639 cm^-1^ were remarkably higher in the vegetative cells, indicating that these peaks are potential differentiation markers for *Anabaena*.

We next plotted the normalized band intensities at 1629 and 1639 cm^-1^ against the “distance” to see the variation of phycocyanin and allophycocyanin in the individual cells (Figures F*–*K). Here, the “distance” represents the number of vegetative cells from a nearby heterocyst (Figure B). The ranges of the former and latter band intensities among the vegetative cells were 0.0039∼0.0079 and 0.0040∼0.0076, respectively. Especially, the upper limit of the band intensity among the heterocysts was lower than the lower limit among the vegetative cells in any filament (Figures F–K). That is, all the vegetative cells indicated the unique higher band intensities of phycocyanin and allophycocyanin. However, we found that the band intensities at 1629 and 1639 cm^-1^ were decreased when vegetative cells were located far from a nearby heterocyst. In the case of the Filament 1, these band intensities were statistically decreased when the “distance” was more than 9, as compared with the intensities in neighboring cells of heterocyst (*p*<0.05, Welch’s *t*-test, Figures F and I). In the case of the Filaments 2 and 3, the band intensities of phycocyanin and allophycocyanin were statistically decreased when the “distance” was more than 10, as compared with the intensities in neighboring cells of heterocyst (*p*<0.05, Welch’s *t*-test, Figures G, J, H, and K).

On the other hands, we also plotted the normalized band intensity at 1328 cm^-1^ to see the variation of chlorophyll *a* in the same manner (Figures L*–*N). The range of band intensities at 1328 cm^-1^ was 0.0021∼0.0059 among the vegetative cells. As Figures L–N show, the band intensities at 1328 cm^-1^ were not statistically decreased when vegetative cells were located far from a nearby heterocyst (*p*>0.05, Welch’s *t*-test). In any filament, the band intensities at 1328 cm^-1^ were statistically identical without regard to the “distance” (*p*>0.05, Welch’s *t*-test, Figures L–N). Thus, we considered that phycocyanin and allophycocyanin were already began to be decomposed in vegetative cells, which were located far from a nearby heterocyst.

As Figures F–H show, some vegetative cells (cells *a–g* in Figures F–H) showed specifically lower band intensities at 1629 cm^-1^. The intensities at 1629 cm^-1^ in cells *a–g* were greater than twice those of the s.d., which was calculated from the intensities at 1629 cm^-1^ in all vegetative cells per each filament. We found that all of them were located in the intermedium region between two existing heterocysts. Moreover, the same vegetative cells *a–g* also showed specifically lower band intensities at 1639 cm^-1^ (Figures I–K).

Given that the band intensities at 1328 cm^-1^ in the vegetative cells *a–g* were not specifically lower (Figures L–N), we considered that the specific decomposition of phycobilisome was occurred preferentially. In fact, we confirmed that the cell *c* was differentiated into a heterocyst 8 hours later after the Raman spectral measurement. Thus, we considered the cell *c* was a “pro-heterocyst” cell when the Raman spectrum was measured. The “pro-heterocyst” is known to start to decrease phycobilisome, and it is still a vegetative cell functionally and morphologically [8]. Although we did not observe the differentiation of the other cells until 8 hours later after the Raman spectral measurement, we were able to observe the specific decomposition of the phycobilisome before the initiation of a cellular expansion by using a Raman microscope.

Our result suggests that the amount of phycocyanin and allophycocyanin was not decreased when the vegetative cell was closed to heterocysts. This is because nitrogen compound (ammonium ion) is supplied sufficiently to vegetative cells from a nearby heterocyst. However, when the cells are deficient of the nitrogen compound, it is known that vegetative cells initiate the decomposition of phycobilisome to gain ammonium ion [8, 9]. In this study, we observed that the vegetative cells, which were located far from a nearby heterocyst, initiated the decomposition of phycobilisome before the differentiation. In fact, we confirmed that one of the cells *a–g* was finally differentiated into a heterocyst. Thus, we were able to observe the precede decomposition of the phycobilisome in the vegetative cell, which would be the candidate cells for heterocyst. In future, we can potentially predict which cell will differentiate or not by measuring Raman spectra in individual cells.

## Figure legends

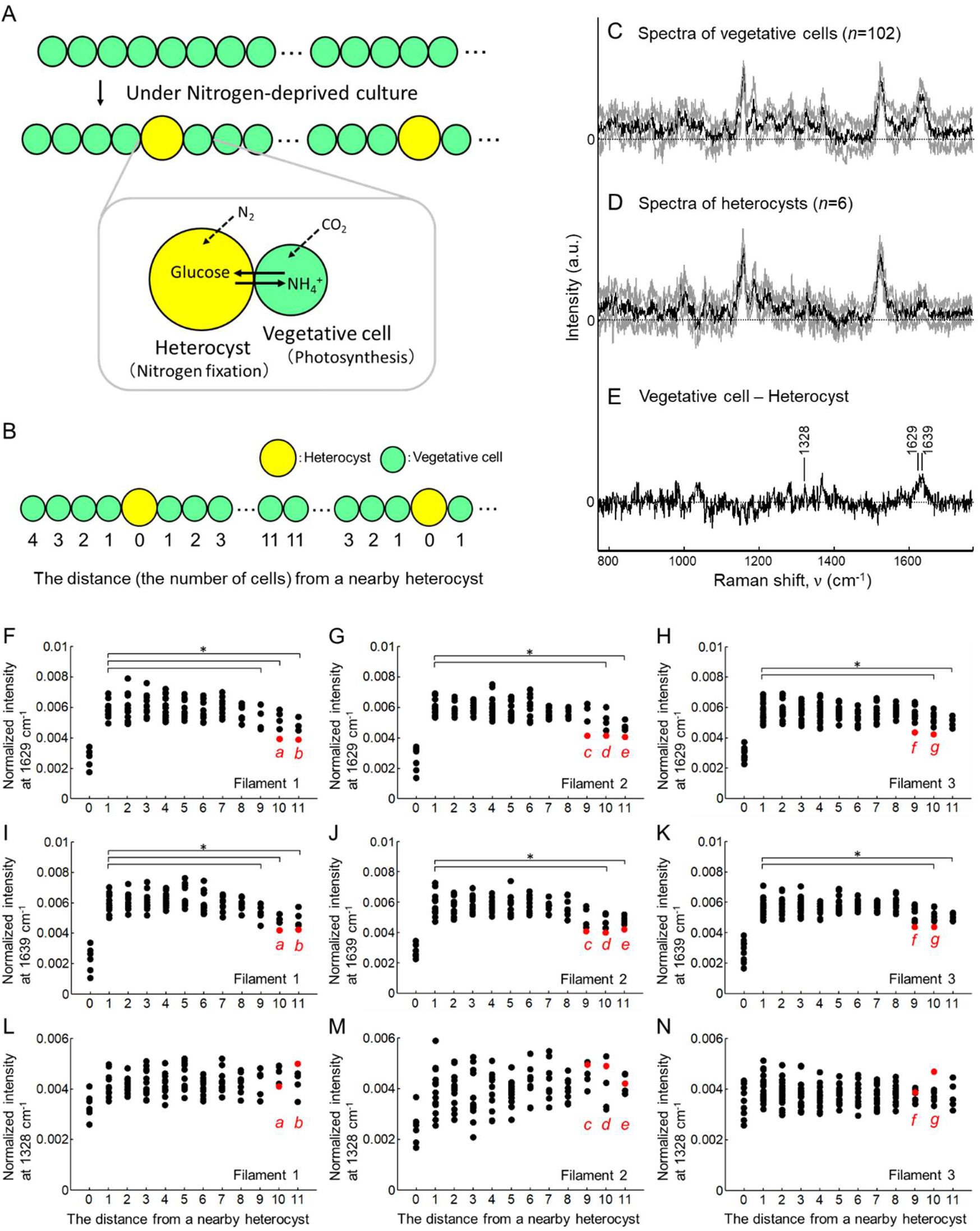
(A) Schematic representation of the heterocyst pattern of *Anabaena* sp. PCC 7120 under nitrogen-deprived conditions. A heterocyst and neighboring vegetative cells exchange the carbohydrates and nitrogen-based compound through periplasm surrounding the cells. (B) Schematic representation of counting the “distance” from a nearby heterocyst. In the case of heterocyst, the distance is counted as 0. Vegetative cells in a filament were numbered along each filament starting from a nearby heterocyst. (C, D) The normalized Raman spectra of the vegetative cells and heterocysts with excitation at 785 nm. Black lines represent the mean spectrum and gray lines represent s.d. We measured the Raman spectra of all vegetative cells and heterocysts in a filament (Filament 1) for every single cell. The Raman bands labeled by arrows (1328, 1629, and 1639 cm^-1^) were representative bands assigned to vibrations of chlorophyll *a*, phycocyanin, and allophycocyanin, respectively. (E) The difference between the Raman spectra (C) and (D). The Raman spectrum in Figure D was subtracted from that in Figure C. (F*–*N) Distributions of the normalized band intensities at 1629, 1639, and 1328 cm^-1^ at different distances from a nearby heterocyst in the three *Anabaena* filaments (Filament 1–3). The number of the data points was 108, 120, and 169, which corresponds to the number of cells in the Filaments 1–3, respectively. The Filaments 1, 2, and 3 included 6, 7, and 10 heterocysts, respectively. The vegetative cells colored in red were the cells *a–g*, which showed the specifically low intensities at 1629 and 1639 cm^-1^ (the intensities were more than twice that of the s.d. in each filament). In Figures F–H, we performed statistical analysis on the Raman band intensities at 1629 cm^-1^ between the cells next to a heterocyst (the “distance” is 1) and other cells (the “distance” is 2∼) in order. In Figures J–K (and Figures L–N), we also performed statistical analysis on the Raman band intensities at 1639 (and 1328) cm^-1^ in the same manner. Statistical difference is indicated by the following symbols. *: *p*<0.05.

## Acknowledgements

We thank Dr. Shin-ichi Morita (Tohoku University), Sota Takanezawa (Nikon), and Hideo Iwasaki (Waseda University) for valuable comments and advice. This work was supported by MEXT KAKENHI (11J06592, 17K15151, 22K15152) to J.I. and MEXT KAKENHI (21K13327) to Y.I.

## Author contributions

JI conceived the project, performed the experiments. JI and YI analyzed the data. JI and YI wrote the manuscript and approved the submitted version.

## Materials and Methods

### Bacterial strains and culture

*Anabaena* sp. PCC 7120 (wild type) were grown in 25 ml of BG-11_0_ (lacking sodium nitrate) liquid medium at 30℃ under illumination with white fluorescent lamps (FL30SW-B, Hitachi co.) at 45 μM photons m^-2^s^-1^. The culture was shook and incubated at 120 rpm until an optimal density at 730 nm (OD_730_) reached about 0.4–0.5. The liquid culture was washed three times by using BG11_0_ liquid medium, diluted to an OD730 of ∼0.2, and underlain beneath a fresh BG-11_0_ solid medium plate containing 1.5 % agar solution (Becton, Dickinson and company, USA) with a bottom dish glass. The sample was placed in a Raman microscope (as mentioned below) kept at 30℃ under illumination with white fluorescent lamps at 45 μM photons m^-2^s^-1^.

### Raman microscope and spectral pre-treatments

In Via confocal Raman spectrometer equipped with a CCD camera (inVia Reflex, Renishaw co.) was used to measure the Raman spectrum. The excitation wavelength was at 785 nm. We measured Raman spectra of individual vegetative cells and heterocysts by selecting the central points of the cells. A typical Raman spectrum of a small confocal volume in the cytoplasm (horizontal diameter, ∼ 1 μm) of a single living vegetative cell (∼ 3 μm diameter) yields a sufficient signal-to-noise ratio for analysis (∼1 s per pixel, with a 785 nm laser at ∼20 mW directed at the confocal volume). In this study, we corrected the baselines of Raman spectra. The baseline-corrected Raman spectrum *y*’(*v*) was calculated as *y’*(*v*) = *y*(*v*) *-y*_poly_(*v*), in which *y*_poly_(*v*) is a fitted polynomial curve constructed by the following procedures. (i) For a spectrum truncated between the minimum Raman shift position *v*_min_ and the maximum position *v*_max_, we selected the degree of the function *d* to fit the baseline using a polynomial function (*d*=3). (ii) Using the least squares method, we first fitted the polynomial function *y*_poly_ to the Raman spectrum *y*. (iii) We divided the Raman spectrum *y* into upper and lower parts, relative to the fitted baseline *y*_poly_. (iv) The number of data points on the upper side of *y* was defined as *N*_*A*_, and the number on the lower side of *y* was defined as *N*_*B*_. If *N*_*A*_ *< N*_*B*_, we removed the upper part of *y* from the whole of *y*, and we replaced the Raman spectrum *y* with the lower part of the spectrum. Then, we repeated the procedure (ii). When *N*_*A*_ *> N*_*B*_, the baseline was considered the best fit and optimal.

